# Hypothetical gene *Rv0495c* regulates redox homeostasis in *Mycobacterium tuberculosis*

**DOI:** 10.1101/2023.08.22.554105

**Authors:** Rahul Pal, Sakshi Talwar, Manitosh Pandey, Vaibhav Nain, Taruna Sharma, Shaifali Tyagi, Vishawjeet Barik, Shweta Chaudhary, Sonu Kumar Gupta, Yashwant Kumar, Ranjan Nanda, Amit Singhal, Amit Kumar Pandey

## Abstract

*Mycobacterium tuberculosis* (Mtb) has evolved sophisticated surveillance mechanisms to regulate and neutralize redox imbalances and associated lethal consequences. Failing this, the accumulated ROS induces toxicity by oxidizing a variety of biological molecules including proteins, nucleic acids and lipids. In the present study we identified Mtb’s *Rv0495c* gene as an important regulator of oxidized cytosolic environment. Compared to wild type Mtb strain lacking *the Rv0495c* gene, Δ*Rv0495c*, had increased ROS and NAD^+^/NADH ratio creating a highly oxidized intracellular environment. Δ*Rv0495c* strain demonstrated slow growth phenotype under *in vitro* and *ex-vivo* growth conditions and demonstrated enhanced susceptibility to drugs, oxidative, nitrosative and hypoxic growth conditions. In addition, the increase in the superoxide radicals triggered a Fenton-like reaction rendering the Δ*Rv0495c* susceptible to free iron. The increase in the intracellular ROS levels of the Δ*Rv0495c* was further corroborated by an increase in the expression of proteins involved in antioxidant defense and enhanced ROS-mediated oxidation and degradation of mycobacterial lipids. This superoxide-induced lipid degradation resulted in altered colony morphology and loss of membrane integrity in the Δ*Rv0495c*. Surprisingly, despite showing a growth defect phenotype in an *ex-vivo* macrophage infection model, the absence of the *Rv0495c* gene in Mtb enhanced the pathogenicity and augmented the ability of the Mtb to grow inside the host. Gene expression analysis revealed a Rv0495c mediated immunomodulation of the host controls inflammation and helps creates a favorable niche for long-term survival of Mtb inside the host. In summary, the current study underscores the fact that the truce in the war between the host and the pathogen favors long-term disease persistence in tuberculosis. We believe targeting Rv0495c could potentially be explored as a strategy to potentiate the current anti-TB regimen.

## 1. Introduction

*Mycobacterium tuberculosis* (Mtb), the etiological agent for tuberculosis (TB), continues to pose a significant global health challenge. This pathogen has evolved mechanisms that allow it to persist and thrive within the human body, evading the immune system and resisting conventional treatments. Factors such as the reactivation in of the infection in the latently infected individuals and emergence of drug resistant strains contribute to the ongoing TB burden worldwide. The emergence of the COVID-19 pandemic has led to setbacks in providing basic TB services and reducing the TB burden. As a consequence, a decline in TB diagnosis and treatment rates has resulted in a significant increase in TB-related deaths which accounts for an estimated 1.3 million deaths worldwide in 2022 due to tuberculosis [1].

Our understanding of host-pathogen interactions, particularly in the context of Mtb infection, remains elusive and continues to be actively pursued by researchers worldwide. Mtb has evolved multiple mechanisms for its survival within the host [2]. Intracellular pathogens like Mtb are challenged with exogenous oxidative stress which is generated by the host as a part of its defence system. The induction of oxidative stress is part of an antimicrobial strategy crucial for host defence against Mtb [3, 4]. Reactive oxygen species (ROS) and nitrogen species (RNS) are generally considered to be mediated through the Fenton reaction[5]. Excess formation of ROS can initiate a series of chemical reactions and cause damage to cellular components, which includes genetic material, proteins and lipids [6, 7]. A minor alteration in the composition of the cell wall lipids in Mtb substantially affects the ability of the pathogen to hide, survive and persist all critical to establish a chronic infection [8]. These lipids not only maintain the integrity of the cell wall but also modulate the immune response of the host during infection [9]. Crucial components of the mycobacterial cell wall, which are required for bacterial virulence includes mycolic acids (MA), phthiocerol dimycocerosates (PDIM) and lipoarabinomannan (LAM) [10]. The ability of the pathogen to survive inside the host is supported and regulated by different Mtb proteins. Since, a quarter of the Mtb genome encoded proteins are yet to be characterized, we hypothesize that a majority of these pathways are unknown and needs to be explored. Some of these hypothetical genes have been speculated to encode for proteases [11], lipases, translocases and esterases[12].

Rv0495c is one such uncharacterized hypothetical protein encoded by the gene *Rv0495c*. This gene is non-essential and is reported to be conditionally essential for the growth on cholesterol [13]. In this study, we showed that *Rv0495c* is crucial for maintaining intracellular redox homeostasis. We observed that the Mtb strain lacking the *Rv0495c* gene (Δ*Rv0495c*) demonstrated a slow-growing phenotype and was hypersensitive to all first-line anti-TB drugs. In addition, relative to the wild type, Δ*Rv0495c* was found to be more sensitive to the oxidative, nitrosative, membrane and hypoxic stress conditions. We further demonstrated that a relative increase in the cytoplasmic ROS enhanced oxidative degradation of mycobacterial lipids in Δ*Rv0495c*. As a consequence, Δ*Rv0495c* demonstrated an altered colony morphology and a loss of membrane integrity. Finally, despite showing a defect in its ability to grow inside the macrophages, the absence of the *Rv0495c* gene augmented the ability of the Mtb to grow in an *in-vivo* mice model reflected in terms of high bacterial load and enhanced pathogenicity.

## 2. Material and methods

### 2.1. Generation of mutant and confirmation

A null mutant of *Rv0495c* was derived from *M. tuberculosis* H37Rv parental strain using homologous recombination between allelic exchange substrate (AES) and mycobacterial genome [14]. Briefly, 1000bp flanking regions of the target gene, *Rv0495c*, were cloned into pJM1 suicidal vector and electroporated in H37Rv competent cells using a standard protocols [15] and transformants were selected over 7H11 plates supplemented with Hygromycin. The mutant strain was screened for the loss of gene due to recombineering using qPCR (Figure 1A, B and C). To complement the *Rv0495c* mutant, the *loxP*-flanked chromosomal hygromycin-resistance gene was excised out by the expression of Cre recombinase. The unmarked null mutant strain thus obtained was transformed with an expression plasmid pJEB402 harboring the *Rv0495c* gene. Strains and primers used in this study were listed in supplementary information Table S1 and Table S2.

**Figure 1.**
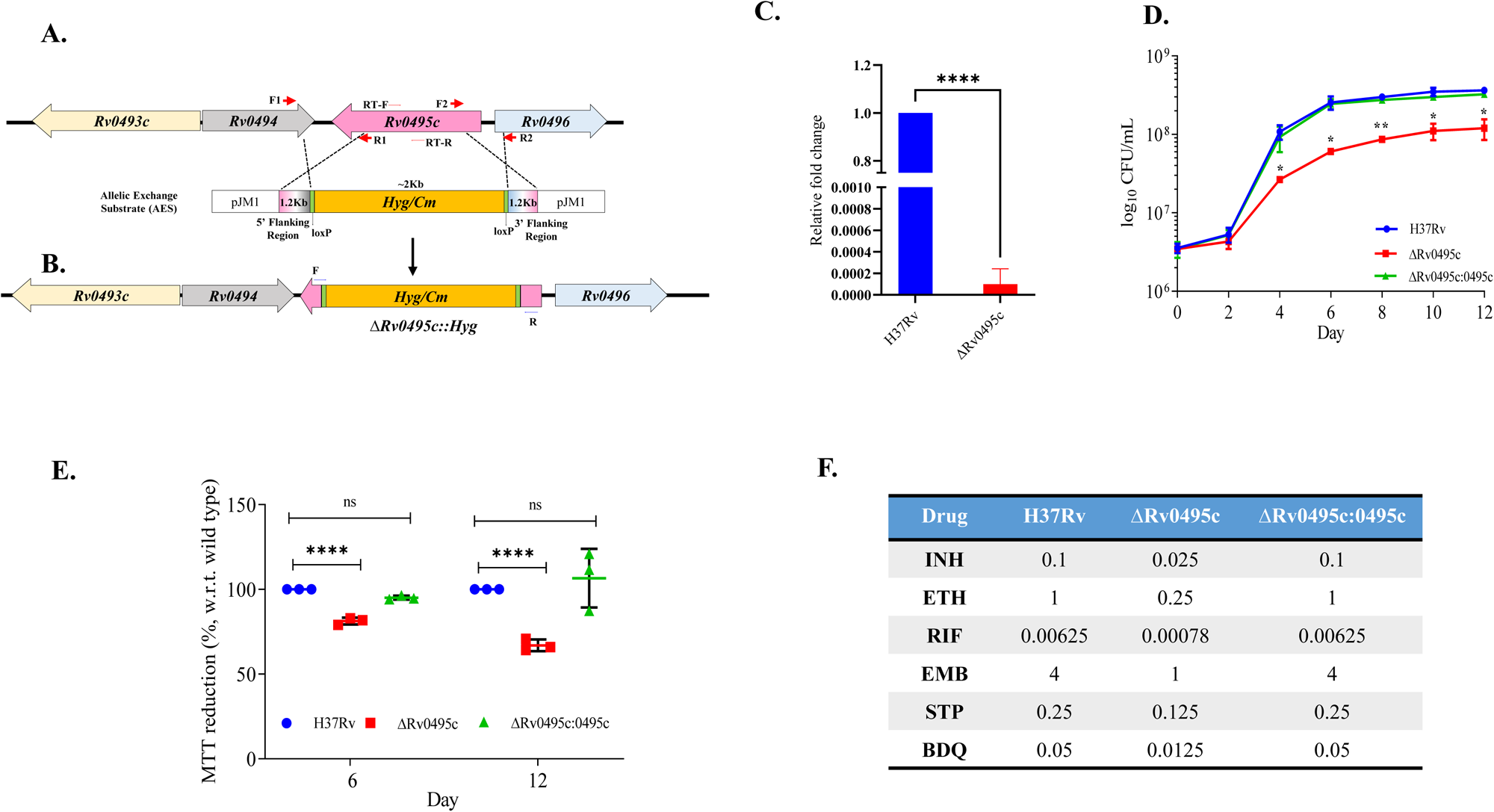
**Generation of Mtb *Rv0495c* deletion mutant and its growth characterization. (A**) Schematic representation of the homologous recombination between the upstream and downstream region of *Rv0495c* gene cloned in the pJM1 suicidal vector and H37Rv genome. (**B**) *Rv0495c* gene replaced by the hygromycin cassette in the H37Rv genome due to homologous recombination, generating a deletion mutant. (**C**) Confirmation of mutant *ΔRv0495c* was done using RT-qPCR as described in material and methods section. **(D)** Growth curve of H37Rv, *ΔRv0495c* and *ΔRv0495c:0495c* strains grown in 7H9 medium enriched with 10% OADS and 0.05% Tween 80 accessed by CFU on 7H11 medium enriched with 10% OADS, colonies were counted after visible growth. **(E).** measurement of metabolic activity of *Mtb* strains using resazurin salt color change from blue to pink indicates reduction of NADH to NAD^+^ levels. **(F)**. MIC was determined by MABA (microplate alamar blue assay) method, performed in the presence of anti-tb drugs Isoniazid (INH), Ethionamide (ETH), Rifampicin (RIF), Ethambutol (EMB), Streptomycin (STP), Bedaquiline (BDQ). Color change from blue to pink/or/purple depicts the growth, each drug was confirmed in at least duplicates.

### 2.2. *In-vitro* growth assessment

To determine growth-related defects due to the deletion of *Rv0495c*, bacterial strains were grown in 7H9 broth. The viable bacterial count was determined by CFU plating at different time points on 7H11 agar plates. The log phase cultures of H37Rv, *ΔRv0495c* and *ΔRv0495c:0495c* strains were washed with PBS twice. Cells were then inoculated into 7H9 broth at an OD of 0.05. The aliquots of the cultures were taken at different time points and plated on 7H11 plates supplemented with 10% OADS for bacterial growth evaluation[16].

### 2.3. Measurement of metabolic activity

The metabolic activity of Mtb strains was estimated using PrestoBlue^TM^. Resazurin color conversion is based on the reduction of the non-fluorescent form of the dye (blue) to an oxidized fluorescent form (red-purple). The biochemical conversion of color from blue to red is a direct measurement of the metabolic activity of live cells. To estimate the metabolic activity of the mycobacterial strains H37Rv, *ΔRv0495c* and *ΔRv0495c:0495c* were grown in 7H9 broth. Cells were washed with PBS and OD was equalized to 0.4-0.5. 200µl cells were then aliquoted in 96 well-plate and 20µL of PrestoBlue reagent (Life Technologies) was added followed by incubation overnight at 37°C. Photographs were captured on the following day.

### 2.4. NADH:NAD^+^ estimation Assay

To measure NADH and NAD^+^ in the mycobacterial strains H37Rv, *ΔRv0495c,* and *ΔRv0495c:0495c* were grown in 7H9 broth. At indicated time points, cells equvalent to 0.5 OD were harvested, washed and resuspended in PBS buffer before adding the MTT [3-(4,5-dimethylthiazol-2-yl)-2,5-diphenyl tetrazolium bromide, 4.2 mM] (Himedia) as directed by manufactures kit. Cells were then incubated for 30 mins at 37°C. NADH and NAD^+^ concentrations were obtained by spectrophotometrically measuring the rate of MTT reduction recorded by taking OD at 570 nm [17]. The concentration of nucleotide (NADH/NAD^+^) is proportional to the rate of MTT reduction.

### 2.5. Colony morphology

10µl of culture (OD_600_:0.1) of the various Mtb strains (*H37Rv, ΔRv0495c and ΔRv0495c:0495c*) were spotted on 7H11 agar plates followed by incubation of the plates at 37^°^C for 12-15 days. The images of growth were captured using the camera.

### 2.6. Drug susceptibility assay

MIC for the anti-TB drugs (RIF, INH, EMB, STP, PYZ) was determined as described previously[18]. Briefly, bacterial cells were grown to mid log phase OD 0.4 and washed with PBS. Anti-TB drugs were serially diluted (2-fold) in 96 well plate. Cells were added at a final density of OD 0.005 to the wells containing drugs the plates were then incubated at 37^°^C for 7 days. 20µl of alamar blue was added to each well of the plates and again incubated for two more days for color development. The color change from blue to pink, by the reduction of resazurin was considered as the presence of bacterial growth. MIC was defined as the lowest concentration of the drug that inhibited bacterial growth.

### 2.7. Stress experiments

Wild type *H37Rv, ΔRv0495c* and *ΔRv0495c:0495c* strains were grown in Middlebrook 7H9 medium. Mid-log-phase cultures were washed with 7H9 medium and exposed to 50µM CHP (cumin hydroperoxide) for 6 hours to generate ROS and 200 µM NO adduct, DETA-NO (RNS) for 48 hours. DETA-NO was replenished after 24 hours. To examine the cell membrane integrity of the strains, the cultures were exposed to 0.1% SDS (surfactant). Culture aliquots were taken at different time points and plated on 7H11 plates supplemented with 10% OADS for the viable bacterial count. For i*n-vitro* hypoxia, bacterial strains were grown in screw cap vials filled 3/4^th^ with bacterial suspensions at an of OD 0.1. Methylene blue (0.01%) was added as an indicator for the development of hypoxia which decolorized at oxygen concentrations of 1.0% or less. Cells were taken at different time points, serially diluted and plated on 7H11 agar plates for growth assessment[19].

### 2.8. RNA extraction and real-time PCR

RNA was extracted using the Trizol method followed by DNase treatment. The quality of the RNA was determined spectrophotometrically by measuring the 260/280nm and 260/230nm ratios. The cDNA was synthesized using an Superscript-IV RT-(Invitrogen) according to the manufacturer’s protocol. The qPCR was set up using TB Green (SYBR) Premix, in QuantStudio7 Pro real-time PCR system (Applied Biosystems). *SigA* was used as an internal control.

### 2.9. Mycobacterial RNA Sequencing and analysis

RNA of different strains was examined using Agilent Bioanalyzer and RNA quality was assessed using a RIN number range of 5.8 to 9.8. 500ng of RNA was used to eliminate rRNA using RiboMinus bacterial transcriptome isolation kit. Using the Lexogen SENSE whole RNA-seq Library Prep kit, cDNA libraries were generated from the rRNA-depleted RNA. A Perkin-Elmer Labchip equipped with a DNA high-sensitivity reagent kit was used to analyze the length distribution of the cDNA libraries. On an Illumina HiSeq 2000 system (Illumina, San Diego, CA; 16 samples/lane), all samples underwent an indexed paired-end sequencing run of 2 ×51 cycles. Applying bowtie2’s default settings, raw reads (FASTQ files) were mapped to the *M. tuberculosis* H37Rv. The genomic annotations from the GenBank file were used to count the genes using feature Counts, a tool included in the Subread suite. Differential gene expression assessment was performed using DESeq2 and the gene counts. Benjamini and Hochberg’s approach was used during multiple testing corrections. Statistical significance was determined by *P-value < 0.05*. The statistical programming language R, version 3.3.1, was used for analyses.

### 2.10. Intracellular redox measurement

To measure intracellular redox levels in mycobacterial strains we used a genetic biosensor (Mrx1-roGFP2) [20]. *Wild type* Mtb and *ΔRv0495c* expressing Mrx1-roGFP2 were treated with 50µM of CHP as an experimental control. After washing three times with PBS, the spectrofluorometric signals were determined for each group by exciting biosensor at 405nm and 490nm while recording fluorescence emission at 510nm using Multimode Spectrophotometer (SpectraMax M2e, Molecular Devices). The ratio of fluorescence at 405/510nm and 490/510nm was calculated and compared for both wild-type and mutant strains to determine the redox state of bacterial cytosol.

### 2.11. Cellular ROS Assay

ROS production was also assessed using a fluorescent dye, 5-(and 6)-chlor-omethyl-20, 70-dichlorodihydrofluorescein diacetate, acetyl ester (CM-H_2_DCFDA; Invitrogen USA, Thermo Fisher Scientific). Mtb cells are grown in Middlebrook 7H9 broth till the OD reaches 0.5-0.6. Cells were then aliquoted and incubated with 20 mM of CM-H_2_DCFDA in the dark (30 min) at 37°C. Cells were fixed with 4% PFA for 20 minutes and then washed once with PBS to remove the excess dye. After washing analyzed by FACS Canto Flow Cytometer (BD Biosciences, San Jose, CA). CM-H2DCFDA fluorescence was determined (excitation wavelength, 488 nm and emission wavelength, 530 nm) by measuring 10,000 events/sample [21]. CHP was used as a positive control and an antioxidant ascorbic acid (AA) as a negative control.

### 2.12. Intracellular metal ions estimation

Mycobacterial strains *H37Rv, ΔRv0495c, and ΔRv0495c:0495c* were grown in 7H9 broth. For intracellular Fe estimation, bacterial cells were harvested on the 6^th^ and 12^th^ day, washed and resuspended cells in deionized water (ICPMS-grade) PBS buffer followed by addition of 200 µl of 70 %conc. HNO_3_ (Trace metal analysis-LCMS grade). After transferring the acid-digested cells to sample vials (MG5, Anton Paar, USA), 100 µl of 30% H_2_O_2_ was added. The vials were then subjected to microwave digestion with a ramp of 250W for 10 minutes and power hold of 250W for 10 minutes. Digested sample were then diluted (1:100) using de-ionized water [22, 23]. Next, these diluted samples were analyzed through ICP-MS (iCAP TQ ICP-MS, Thermo Fischer Scientific). 57Fe standard was run at various concentration (1ppb to 1000ppb) to prepare the calibration plot. Briefly the samples were introduced to micromist nebulizer with the help of sample capillary. The nebulizer supply pressure was set at ≈ 3 bar and the peristaltic pump used to aspirate sample via sample capillary was revolving clockwise at the speed of 40 rpm. The aerosols generated through the nebulizer were ionized with plasma by passing through the sample cone followed with skimmer cone. The instrument exhaust was maintained between 0.4 to 0.7 mbar. Samples were run with a dwell time of 0.1 sec and 1% HNO_3_ was run for 30 sec between samples to remove carryover from previous sample. Average value of three runs was used to calculate iron concentration in each sample. All the samples were run in KED mode (Kinetic energy dissociation) to remove polyatomic interference. Qtegra software from Thermo scientific was used to operate the instrument. ICP-grade water was used for standard preparation, sample dilution and as blank. Data obtained were then normalized with the the total bacterial count estimated by CFU plating.

### 2.13. Ethidium bromide (EtBr) uptake

The uptake of EtBr was determined as previously described with some modifications[24]. Briefly, cells were grown to an OD of 0.6-0.8, and washed with PBS. OD of each culture suspension was adjusted to 0.4 in PBS, transferred to 96-well plate followed by the addition of 5 μM of EtBr. Finally, fluorescence was measured using a Multimode Spectrophotometer (SpectraMax M2e, Molecular Devices). Fluorescence was measured at the excitation wavelength of 530 nm and an emission wavelength of 590 nm. Culture suspensions without EtBr were used as a control for the normalization of auto-fluorescence.

### 2.14. TEM analysis

Specimens for transmission electron microscopy were prepared as previously described[25]. Briefly, an aliquot of cells grown in 7H9 broth (without Tween 80) containing approximately 5×10^7^ cells was pelleted by centrifugation and fixed with a combination of 2.5% paraformaldehyde along with 2.5% glutaraldehyde. Electron microscopy was performed with a JEM1400 Flash Transmission Electron Microscope operating at 120 kV (ATPC, Regional Center for Biotechnology, India).

### 2.15. Lipid extraction and analysis

Wild-type H37Rv, *ΔRv0495c,* and *ΔRv0495c:0495c* strains were grown in 10 ml of 7H9-enriched media. The cells were harvested and washed with PBS, and resuspended in chloroform-methanol (2:1, vol/vol) mix and left overnight on shaking. This was followed by sequential extraction with chloroform-methanol (1:1, vol/vol) and chloroform-methanol (1:2, vol/vol) [26]. For lipid analysis 10 μl of each lipid was spotted onto a pre-dried TLC plate (Silica gel; Sigma) at a distance of 2cm upward from the end of the plate. The solvent systems used for lipid analysis are listed in (Supplementery Information 4, Table S3). Development of TLC was done using iodine vapors.

### 2.16. Mass spectrometry analysis

The total cellular extracted lipids (as described above) were separated using Ultra-High-Performance Liquid Chromatography (UHPLC) systems coupled with orbitrap fusion mass spectrometry. Briefly, an ACQUITY HSS T3 (2.1 mm × 100 mm × 1.8 μm, Waters) column with a sample injection volume of 5 μL was employed for the separation of lipids. The column temperature was maintained at 40 °C throughout the analysis [27]. Solvent system-A was water: acetonitrile in a 40:60 vol/vol ratio, and solvent system -B was 2-propanol: acetonitrile in a 90:10 vol/vol ratio at a flow rate of 0.3 mL/min. The gradient elution started at 30–97 % B for 1–12 min and was maintained for another 3 min, further 15.2–18 min was maintained at 30 % B. Mass spectrometric analysis was performed on a high-resolution mass spectrometer, orbitrap fusion (Thermo Scientific, USA), fitted with a heated electrospray ionization source (H-ESI). The ionization voltage was set at 3000 kV for positive and 3500 kV for negative ion modes. The H-ESI sheath and auxiliary gases were maintained at 60 arbs and 20 arbs, respectively. The temperature of the vaporizer was 310 °C, and the capillary temperature was at 300 °C. The data acquisition was carried out at 120 k resolution for every MS run in a mass range of 200–1200 with an automatic gain control (AGC) target of 200,000 ions, whereas MS/MS resolution was set at 30 k resolution with an AGC target of 50,000 ions. Data obtained were then analyzed using MS-LAMP software using a window range of 0.5 in [M-H] negative ion mode[28] and Mtb LipidDB[29].

### 2.18. Lipid hydroperoxidation assay

The lipid hydroperoxides were determined as previously described [30]. Briefly, the cell pellet (correnponding to 1 OD cells) washed with deionized water twice. The resulting pellet was then dissolved in 1 ml methanol/chloroform (2:1 v/v) solution. The suspension was vortexed at room temperature. From this, 200 µl aliquot was taken and mixed with 800 µl of FOX-II reagents and incubated for 30 min. The level of lipid peroxide in the cell was monitored by measuring the absorbance at 560 nm.

### 2.19. THP-1 infection

The contribution of the Rv0495c protein to help Mtb’s ability to survive inside macrophages was estimated through *in-vitro* infection studies using THP-1 monocyte derived macrophages. Briefly, THP-1 monocytes were seeded into 24-well plates at a density of 5×10^5^ cells/well. Cells were treated with PMA (50ng/ml) for 24 hours to allow them to differentiate into macrophages and adhere to the surface. This was followed by resting for 24 hours. Media was changed 24 hours prior to the infection. Macrophages were then infected with respective strains at an MOI of 1 for 4 h at 37°C and 5% CO_2_ followed by extensive washes with warm PBS to remove extracellular bacteria. Intracellular bacterial count was determined by lysing the cells with 0.01% Triton X-100 at the indicated time points and plating dilutions on 7H11 agar [16].

### 2.20. Animal infection Study

The animal experiment protocol was reviewed and approved by the Institutional Animal Ethics Committee of Translational Health Science and Technology Institute, Faridabad, India (IAEC/THSTI/183), and (IAEC/THSTI/167). Animal experiments were performed in accordance with guidelines provided by the Committee for the Purpose of Control and Supervision of Experiments on Animals (Govt. of India). Pathogen-free C57BL/6 female mice were obtained from the Small Animal Facility, THSTI, India. Eight-week-old C57BL/6 female mice were infected through aerosol exposure using an aerosol generator (Glas-Col) with approximately 100 CFUs of respective strains (*H37Rv, ΔRv0495c* and *ΔRv0495c:0495c*). Mice from each group (n = 5) were sacrificed at different time points (Week 0-, 4-, 8- and 12-weeks post-infection). Their lungs and spleen were harvested and the serial dilutions of the organ homogenates were plated on 7H11 agar plates for bacterial enumeration.

### 2.21. Host RNA Sequencing

NEB Ultra II directional RNA-Seq Library Prep kit protocol was used to prepare libraries for mRNA sequencing (NEB). An initial concentration of 500ng of total RNA was taken for the assay. mRNA molecules are captured using magnetic Oligo d(T)beads (NEB). Following purification, the enriched mRNA was fragmented using divalent cations under elevated temperatures. The cleaved RNA fragments were copied into first-strand cDNA using reverse transcriptase. Second-strand cDNA synthesis was performed using DNA Polymerase I and RNase H enzyme. The cDNA fragments were then subjected to a series of enzymatic steps that repair the ends, tails the 3’ end with a single ‘A’ base, followed by ligation of the adapters. The adapter-ligated products were then purified and enriched using the following thermal conditions: initial denaturation of 98°C for 30sec; 12 cycles of - 98°C for 10sec, 65°C for 75sec; final extension of 65°C for 5mins. PCR products are then purified and checked for fragment size distribution on Fragment Analyzer using HS NGS Fragment Kit (1-6000bp) (Agilent).

### 2.22. Data Availability

All data generated or analyzed during this study are included either in this article or in the supplementary information files. Host RNA sequencing Accession number: PRJEB4000.

### 2.23. Ethics Statement

The animal study was reviewed and approved by the Institutional Animal Ethics Committee (IAEC), THSTI.

### 2.24. Statistical analysis

Data were expressed as mean ± SD, Student’s *t-test* and one-way analysis of variance (ANOVA), was used to determine statistical differences between the groups. GraphPad Prism 9.0 software (GraphPad Software) was used for statistical analysis. A *p-value* of *<0.05* is considered to be indicative of a statistically significant result. *, *p < 0.05; **, p < 0.01; ***, P < 0.001*.

## 3. Results

### 3.1. Absence of *Rv0495c* altered growth, colony morphology and drug susceptibility in *Mtb*

Δ*Rv0495c* knockout strain in Mtb was constructed by inserting a hygromycin cassette in the *Rv0495c* open reading frame (ORF) by homologous recombination (Figure 1A). The insertional inactivation of the *Rv0495c* gene in the double crossover mutant strain was confirmed by amplifying the flanking regions (Figure 1B) and by RT-PCR (Figure 1C). Growth assessment on an enriched media demonstrated a growth defect in mutant strain relative to the wild type, which was completely restored in the complemented strain (Figure 1D). Since the NADH/NAD^+^ ratio is the predominant indicator of the metabolic activity, we estimated the NADH/NAD^+^ ratio in different strains to evaluate their metabolic state. For the estimation of NADH levels we used a resazurin-based assay where, depending on the metabolic activity, the active component is reduced to form either a purple to a pink color product. While pink color developed in the case of the wild type and the complemented strain indicated normal metabolic activity, a purple color observed in the case of the *ΔRv0495c* suggests a reduction in the metabolic activity (Figure S1). The same was further validated using MTT dye-based assay. Low levels of tetrazolium reduction in Δ*Rv0495c* relative to the wild type indicated decreased intracellular levels of NADH/NAD^+^ which is indicative of reduced metabolic activity in the mutant strain (Figure 1E).

Further, in order to investigate the effect of gene deletion on the drug susceptibility pattern, these strains were exposed to various 1^st^ and 2^nd^ line anti-tubercular drugs. Interestingly, regardless of the drugs tested, in comparison to the wild type and the complemented strains, *ΔRv0495c* was found to be 2- to 4-fold more sensitive to various anti-TB drugs tested (Figure 1F). Collectively, these results indicate that the *Rv0495c* gene modulates growth, metabolism and regulates the sensitivity of *Mtb* towards anti-tubercular drugs.

### 3.2. Rv0495c protein helps Mtb survive under various host-induced stress conditions

Growth modulation and altered cell wall permeability suggest that the Rv0495c protein may affect the survival of the pathogen inside the host. Accordingly, we studied the effect of *Rv0495c* gene disruption on mycobacterial survival upon exposure to physiological stressors, such as CHP (oxidative stress), SDS for cell membrane stress and Deta-NO for nitrosative stress. Survival was assessed in terms of colony forming unit (CFU) post-exposure to the stressors. We observed that, relative to the wild type, Δ*Rv0495c* was more sensitive to oxidative (2-fold), membrane (2-fold) and nitrosative stress (1.5-fold) (Figure 2A, B&C). Interestingly, relative to the wild type, Δ*Rv0495c* was also found to be defective in growth under hypoxic conditions. Briefly, the cultures were grown in sealed vials. The utilization of the oxygen present in the headspace eventually creates a low oxygen growth condition. There was a significant mutant-specific decrease in the CFU observed at all the estimated time points i.e., 20-, 40- and 60-days post inoculation (Figure 2D). Overall, the data suggest that the absence of *Rv0495c* renders *Mtb* prone to redox, cell membrane and hypoxic stress conditions. Mtb is known to grow and replicate inside macrophages under conditions that include exposure to high ROS and RNI. To check if Rv0495c has any role in the intracellular survival of the pathogen we used a human THP-1 derived monocyte cell line. Briefly, the THP-1 cells were infected with different Mtb strains and the uptake and the growth were measured by performing CFU assay. The differences in the uptake observed for wild-type, mutant and complementary bacterial strains by THP-1 cells were not significant at 6.0%, 5.3% and 3.6% respectively. Surprisingly, *ΔRv0495c* failed to replicate inside the macrophages and at 7 days post-infection demonstrated a 90% reduction in the growth relative to the wild type and the complemented strain (Figure 2E).

**Figure 2.**
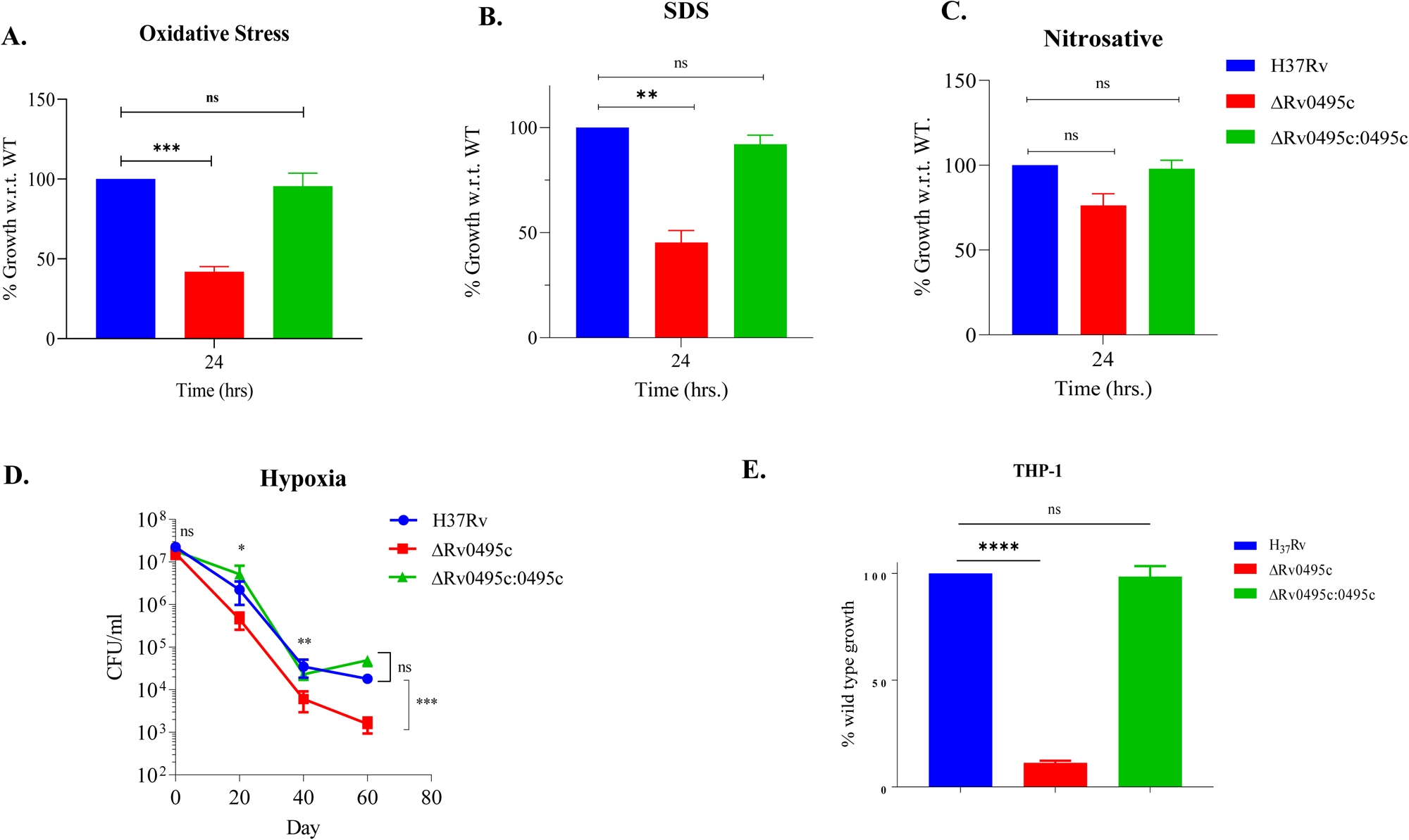
**Effect of host-induced stresses on the endurance of *Rv0495c* deletion mutant**. **A**. 100μM CHP was added to log-phase cultures of *H37RvWT, ΔRv0495c and ΔRv0495c:0495c* strains and growth was monitored after 6 hours of plating data obtained was then normalized to 0 hour and plotted with respect to wild type H37Rv (100%) on the y-axis. **B**. For membrane stress (0.1% SDS for 24 hours) data obtained was then normalized to 0 hour and plotted with respect to wild type *H37Rv* (100%) on the y-axis. **C.** Nitrosative stress (200 μM DETA-NO for 48 hours data obtained was then normalized to 0 hour and plotted with respect to wild type *H37Rv* (100%) on the y-axis. **D.** Hypoxia (O_2_ depletion) in *H37Rv, ΔRv0495c and ΔRv0495c:0495c* strains was measured by Wayne model. Approximately 1.5 × 10^8^ (OD = 0.5) mycobacteria were cultured in screw cap tube. At indicated time points, or bacteria were plated for CFU analysis. (**E)** Growth of the *ΔRv0495c* strain relative to the wild-type strain in THP-1 cell line. THP-1 macrophages were infected at an MOI of 1, and a relative growth difference was estimated by counting colony-forming units at 0hr, 4hr and 24hr after plating. Data was normalized with 0 day and plotted with respect to wild type *H37Rv* (100%) on the y-axis. The data depict here is mean ± SEM, representative of two independent experiments performed in replicates. *, *p* < 0.05; **, *p* < 0.005; ***, *p* < 0.0005.

### 3.3. *Rv0495c* is essential for maintaining redox homeostasis in *Mtb*

In order to identify a set of genes affected by the loss of *Rv0495c* in *Mtb*, we analyzed the differentially expressed genes between the wild type and Δ*Rv0495c* cultures, grown in an enriched media, by quantifying the transcripts using RNA sequencing (Figure 3A). We found that the transcript levels of 23 genes were differentially regulated (FDR<0.05) and a majority of them were found to be involved in redox homeostasis (*whiB3, Rv0575c, Rv3741, Rv3742, ahpC* and *fadE5*), lipid metabolism (*desA2, desA3, FadE5, FadD9, Rv3740*), conserved hypothetical proteins (Rv1734c, Rv2030c, Rv3129, Rv1057, Rv3843c) or involved in pro-drug activation (*ethA, ethR*) (Figure 3B and Supplementary spread sheet 1). The above findings indicate the downregulation of genes required for maintaining redox homeostasis. Therefore, a protocol using a genetically encoded redox-sensitive biosensor [31] was used to determine the intracellular redox state of both the wild-type and the mutant strains [31]. Briefly, Mtb*-Rv*:Mrx1-roGFP2 strain was exposed to CHP and DTT and the change in oxidative/reductive ratio was then compared with *ΔRv0495c*:Mrx1-roGFP2 (untreated) strain. The treatment of CHP in parental H37Rv:Mrx1-roGFP2 strain resulted in an increase in the 390/490nm ratio while DTT resulted in a marked decrease in the 390/490nm ratio when compared to untreated H37Rv:Mrx1-roGFP2 strain. Interestingly, in the case of *ΔRv0495c:*Mrx1-roGFP2 we found a significantly higher oxidative state of cytosol similar to the redox change observed in the case of CHP treated H37Rv:Mrx1-roGFP2 (Figure. 3C) suggesting that *ΔRv0495c* is not able to maintain the intracellular redox balance which is critical for the normal functioning of the pathogen. A higher 390/490 ratio indicates the presence of an oxidized form of mycothiol inside the cytosolic environment, which occurs due to the accumulation of ROS. Therefore, ROS generation was biochemically measured using 2′,7′-dichlorofluorescin diacetate (DCFDA), which upon exposure to ROS converts into a fluorescent form [21]. Consistent with the previously observed data, we found that ROS generation inside *ΔRv0495c* was significantly higher than wild type (Figure S2). Further, high intracellular ROS resulted in an increase in the total Fe concentration inside the cell (Figure 3D and S3). This increase in the intracellular Fe levels together with ROS created a Fenton-like reaction that made *ΔRv0495c* sensitive to free Fe. Supplementation of the growth media with several Fe-binding proteins resulted in the restoration of the growth of *ΔRv0495c* to the untreated wild-type levels (Figure 3E). Additionally, to confirm whether the above phenotype is also be true in *ex-vivo* condition. We supplemented the cell culture media with iron-binding protein Hb-S and a siderophore chelator (mbt-J). As expected, supplementation of both in the cell culture media relieved growth restrictions phenotypes of the *ΔRv0495c*. We found that the intracellular growth of the *ΔRv0495c* was restored to the wild type levels when supplemented with Hb-S whereas supplementation with mbt-J, which has the ability to bind and import iron, not only restored the growth but resulted in 2-fold better growth than the wild type (Figure 3F). Overall, the above data suggest that the *Rv0495c* protein of Mtb is essential for maintaining intracellular redox homeostasis by neutralizing the excess of ROS generated in the cytosol of Mtb during infection.

**Figure 3.**
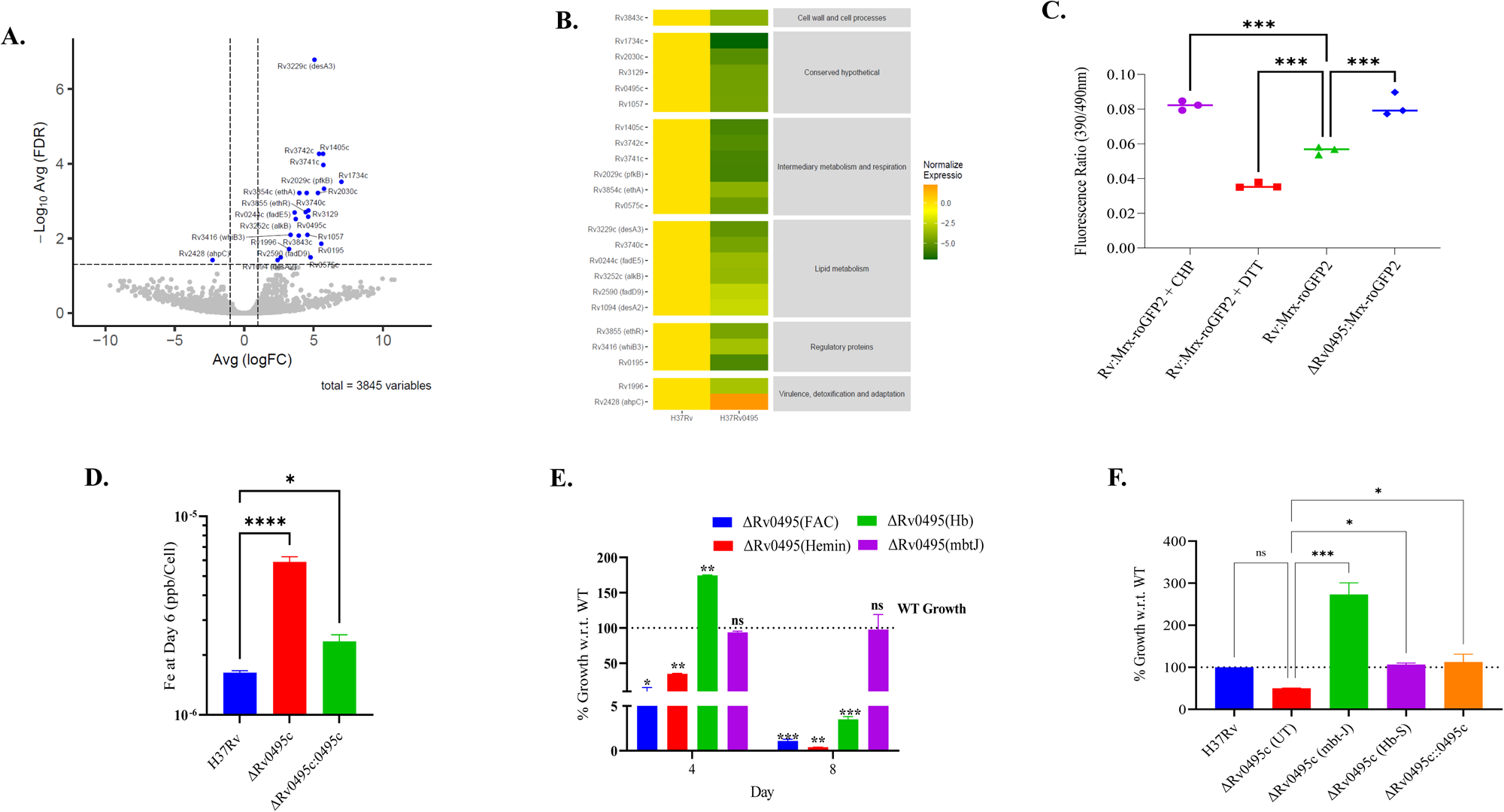
Deletion of *Rv0495c* gene disrupt the capacity of the cell to detoxify the reactive species and alters the intracellular metal ion levels. (A) Volcano plot of all the differentially expressed genes (DEG) tested visualizing fold change in expression and statistical significance. Black dots indicate genes with no significant differences in expression; light blue dots represent genes with *p-value < 0.05*. Whereas, dark blue dots represent genes with false detection rate (FDR) <0.05. The average FDR in expression levels between *H37Rv Vs ΔRv0495c* (*y-axis*) and are plotted against the average log fold changes Avg (Log FC) in expression (*x-axis*). **(B)** Heat map of log fold changes for differentially expressed genes having with FDR <0.05. and clustered according to their category. **(C).** The graph represents the 390/490nm ratios change upon oxidation or reduction of Mrx1-roGFP2 in response to oxidative or reductive stress. Exponentially grown cells of *H37Rv* strains expressing Mrx1-roGFP2 were exposed to with or without 50 μM CHP and 50mM DTT along with ΔRv0495c. Response of ratiometric biosensor was measured at 10 min using spectrofluorometer, as described in (*Materials and Methods)*. Error bars represent SD from the mean. * p < 0.05 (as compared to Mrx*1-Rv-roGFP2*), and (***p < 0.0005 as compared to Mrx1-Rv-roGFP2 treated with DTT), and p>0.05 (ns as compared to Mrx1-Rv-roGFP2 treated with CHP). Data are representative of at least three independent experiments. **(D).** Intracellular iron levels of *H37Rv, ΔRv0495c*, and *ΔRv0495c:0495c* Mtb strain grown in 7H9 medium enriched with 10% OADS and 0.05% Tween 80 was measured at 6^th^ day using ICP-MS. *y-axis* depicts the concentration of iron in ppb/bacteria with respect to time (days) on x-axis. Data represented here was normalized with CFU, and replicated in *n = 3* biological and averages for two independent sets of experiment. **(E). Growth of different iron source.** Growth of *H37Rv, ΔRv0495c*, and *ΔRv0495c:0495c* strains in sautan’s minimal medium supplemented with 10µ ferric ammonium citrate (FAC), 2.5 µM hemoglobin (Hb-B), 10µM Hemin and Mycobactin-J (Mbt-J). Error bars represent SE± of mean values of biological replicates.

### 3.4. Deletion of *Rv0495c* alters the composition of cell membrane-associated lipids

When plated on the agar plates the Δ*Rv0495c* exhibited visibly altered colony morphology possibly indicating a difference in the composition of the cell wall lipid between these strains (Figure S4) which might have altered the cell wall integrity of the *ΔRv0495c*. To evaluate this, we performed the EtBr uptake assay as a measure of membrane permeability in bacteria [32]. EtBr gives fluorescence only upon binding to DNA or cellular components in comparison to its unbound form in an aqueous solution. Interestingly, *ΔRv0495c* showed a higher accumulation of EtBr in contrast to the wild-type bacteria suggesting a more porous and compromised cell wall (Figure 4A). Using Transmission Electron Microscopy (TEM) we observed that in comparison to the H37Rv *ΔRv0495c* bacteria had increased length with a significant decrease in the lipid content in the outer membrane. The colony morphology, stress response, membrane permeability and TEM data (Figure 2B, 4A, 4B, 4C and S4) together suggested that the *Rv0495c* gene regulates the abundance of membrane lipid critical for the cell wall integrity. In order to validate this hypothesis, we analysed the total lipid content of *H37Rv, ΔRv0495c, ΔRv0495c: Rv0495c* strains. We extracted and analysed the total lipids using one-dimensional (1D) -TLC with the solvent system mentioned in (Table S3). As predicted, we found visual differences in the TMM (trehalose monomycolate), methoxy mycolate and the PDIM fraction of the total lipid (Figure 4D, 4E). However, in the case of phospholipid different classes were affected as revealed by 1D and 2D-TLC (Figure S5A and S5B). Further, total cellular lipid extracted were studied using an unbiased approach LC-MS/MS to fully characterize the lipids between the wild type and the *ΔRv0495c* strains. The spectra obtained from LC-MS for each of the samples were processed and analysed for the PCA plot and found significant variations between the lipid contents of H37Rv and Δ*Rv0495c* strains which seems to be resolved in the ΔRv0495c:0495c complemented strain (Figure 4E). To analyse m/z values and identify the lipids, we used MS-LAMP software which works on the basis of MycoMass database. MycoMass categorizes total lipids into six different classes: Fatty acyls (FA), Glycerolipids (GP), Glycerophospholipids (GPL), Prenols (PR), Saccharolipids (SCL) and Polyketides (PK). Under these classified categories, we identified 234 lipids species using MS-LAMP software in [M-H]^-^ Negative ion mode in the 0.5 window range (Supplemnatry spread sheet 2). Here, we focus mainly on the three classes: FA, GP and GPL. Intensities of almost all the sub-classes of lipids were found to be considerably decreased in Δ*Rv0495c* and progressively their levels were restored to wild type strain in the complemented strain. We have recognized a total of 76 FA of which only GMM-MA showed a significant reduction in the *ΔRv0495c* strain. We also observed a significant decrease in the levels of glycerolipids (monoglyceride, diglyceride and triacylglycerides) and triacylglycerides (TG (45:1), TG (46:2) and TG (48:0)). We also attempted to examine phospholipids and found significant differences in 16 subclasses of phospholipids that were less abundant in *ΔRv0495c* strain (Figure 4F). A general reduction in the lipid content across all the categories in the *ΔRv0495c* strain can be very well explained by a significant increase in the lipid peroxidation determined biochemically by FOX-II reagent (Figure 4G). Increased ROS generation leads to peroxidation of cellular lipids which in turn destabilized the cell membrane and renders it sensitive to withstand extracellular assaults [4]. In sum, we report that the absence of *Rv0495c* in Mtb created a highly oxidizing intracellular environment that triggered the peroxidation of the cellular lipids that severely compromises the membrane permeability and cell wall integrity of Mtb.

**Figure 4.**
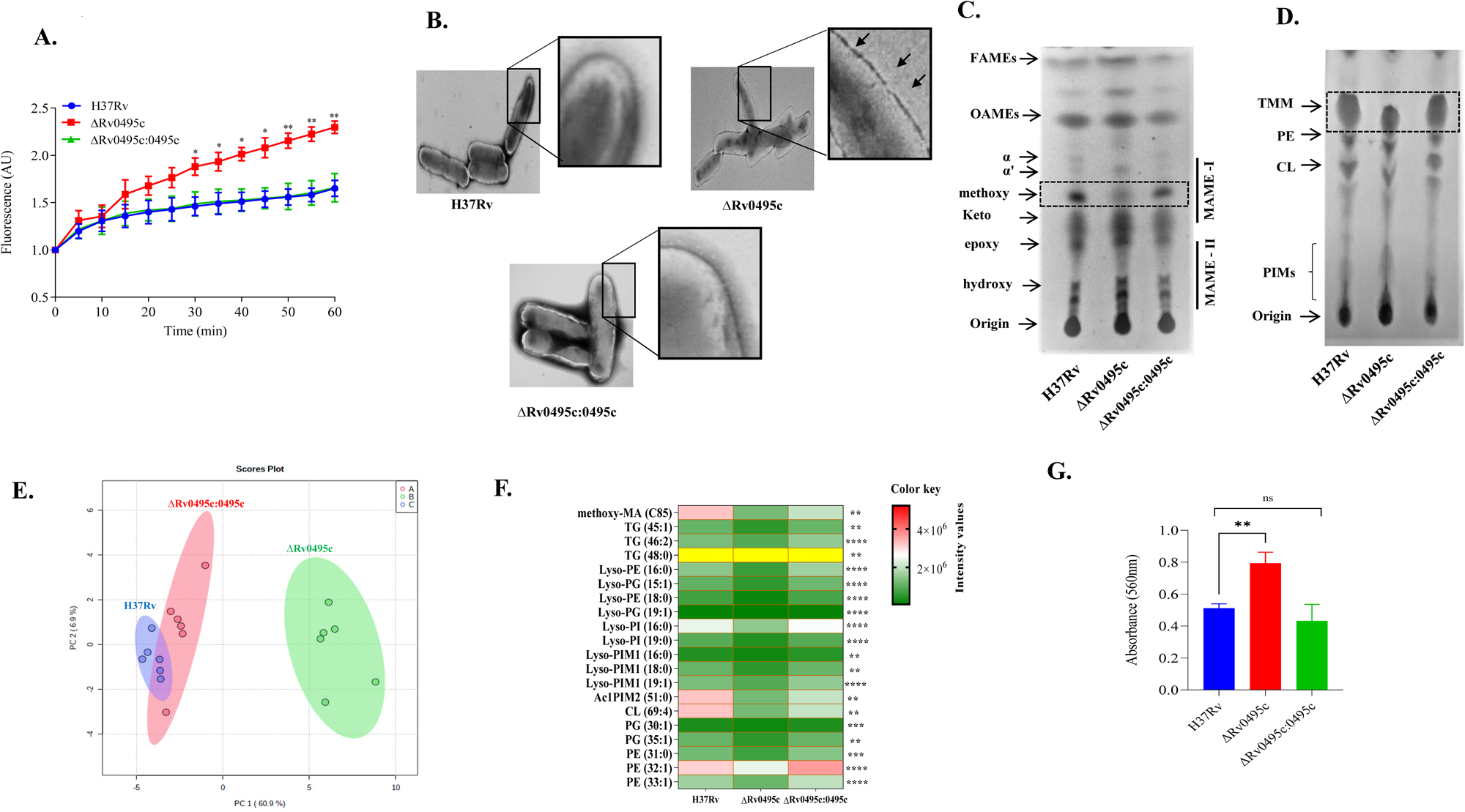
Role of *Rv0495c* in membrane homeostasis. (A) Membrane permeability was accessed by measuring intercellular level of EtBr. Mean of fluorescence ± SD of three independent sets of experiment is depicted on y-axis with respect to time (minutes) on x-axis. **(B)** TEM images showing thin membrane, tampered and elongated morphology of ΔRv0495c as compared to wild type (*H37Rv*), and complimentary strain *(ΔRv0495c:0495c*). **(C)** Lipids were extracted from mid-logarithmic phase cultures of *H37Rv, ΔRv0495c* and *ΔRv0495c:0495c* strains and analyzed for the presence of Mycolic acids were analyzed by1D-TLC using solvent [hexane/ethyl acetate (19:1, v/v)]. For visualization TLC plates were fumed with iodine vapors. **(D)** 1D-TLC using solvent [Chloroform: Methanol: water (65:25:4, v/v/v)] for the analysis of phospholipids (PIMs, PE, CL, TMM, Acylated-PIMs). **(E)** Principal component analysis (PCA) of the identified lipid molecules. PCA plot of the identified lipid molecules from lipidomes of *H37Rv, ΔRv0495c* and *ΔRv0495c:0495c* strains. **(F)** Heat map represents the global lipidomic response of the six biological replicates samples identified in (M-H) ion mode and analyzed using MS-LAMP software in a 0.5 window range. **(G)** Lipid peroxidation in *H37Rv, ΔRv0495c* and *ΔRv0495c:0495c* bar graph represents the mean absorbance at 560nm wavelength.

### 3.5. Δ*Rv0495c* demonstrated an enhanced ability to grow inside the host

We assessed the impact of the deletion of *Rv0495c* on the ability of the pathogen to grow and establish disease in the mice model. Mice were infected with *H37Rv*, *ΔRv0495c* and *ΔRv0495c:0495c* through the aerosol route of infection and estimation of the pathogen load and histopathological analysis was done after harvesting the lungs and spleen 4 and 8 weeks post infection. Surprisingly, high bacillary load was observed in the lungs and the spleen of animals infected with *ΔRv0495c* as compared to H37Rv and *ΔRv0495c:0495c* at both the time points (Figure 5A and 5B). Grossly, the organs isolated from *ΔRv0495c* infected mice had hyperinflammation and had more numbers of granuloma visible on the surface relative to organs isolated from the wild-type infected mice. The histopathological analysis also revealed that the lungs of the mice infected with *ΔRv0495c* had severe pathology and demonstarted high granuloma score relative to the mice infected with wild type Mtb (Figure 5C and 5D). The above data suggests that the absence of *Rv0495c* in Mtb increased the fitness of the pathogen to grow inside the mice. This could also be attributed to the enhanced immunomodulatory properties of the *ΔRv0495c* which increased the ability of the pathogen to grow inside a host.

**Figure 5.**
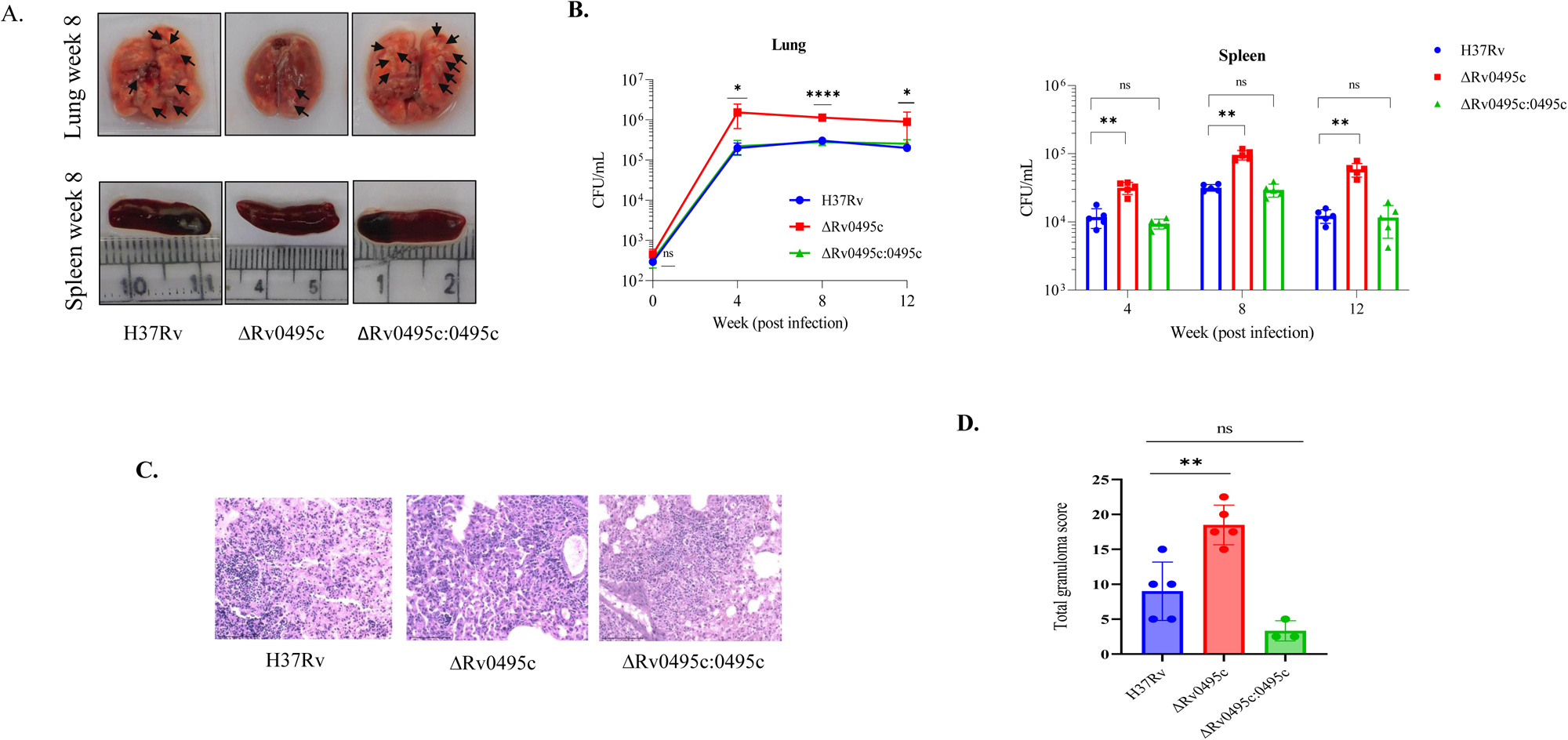
*ΔRv0495c* gene plays a crucial role in the regulated growth of Mtb in the host. **(A)** Assessment of Gross pathology of the lungs and spleens of animals infected with mycobacteria strains at 8 weeks post infection. Images shown here are representative of those obtained after 8 weeks of mice sacrificed **(B)** Bacterial load in mice infected with different Mtb strains through the aerosol route (*H37Rv, ΔRv0495c*, and *ΔRv0495c:0495c*) in the lung and spleen, respectively, at week 0, 4, 8 and 12 post infection. The data represents the average colony-forming count from five animals in each group. Significant differences observed in the groups are marked (ANOVA, *\*P < 0.005*). **(C)** For the assessment of tissue damage lung tissue of mice 8 weeks post infection were stained with hematoxylin and eosin and was photographed at 40X magnification, scale bar 100μm. The images shown are representative of those obtained from 5 animals per time point for each strain. **(D)** bar graph showing the average of granuloma score.

### 3.6. Deletion of Rv0495c alters the host immune response in *M. tuberculosis*

To gain more insights into the specific host factors contributing to the hypervirulent phenotype of the mutant strain, we conducted RNA sequencing on the lung tissues of mice infected with the WT strain and the *Rv0495c* mutant strain. To begin the analysis, we performed a principal component analysis (PCA) to visualize the variations in gene expression profiles between the two strains. The data represented in the first two principal components (PCs) indicated clear separation among both gene clusters (Figure 6A). To investigate the impact of *Rv0495c* on host gene expression, we compared lung tissue gene expression profiles from mice infected with the *Rv0495c* mutant strain and those infected with the WT strain (Figure 6B and 6C). Our analysis revealed a total of 2402 genes differentially expressed between the WT and *Rv0495c* mutant strains. Among the DEGs, 968 were up-regulated, while 1433 were down-regulated in the *ΔRv0495c* relative to the wild type (Supplementary File S3). Figure 6C illustrates the expression levels of the top 20 genes that are upregulated and downregulated in the mutant strain when compared to the WT strain. In the mutant strain compared to the WT strain, the upregulated genes included C1qa (3.4), C1qb (3.2), and C1qc (3.2), which belong to the complement system. The role of the complement system cascade in the pathogenesis of Mtb infection is still unclear, but higher expression of C1q has been associated with more severe clinical conditions. Additionally, IL1a (3.6) and IL1b (5.6), which are pro-inflammatory markers associated with TB, were significantly upregulated in the tissue obtained from mice infected with the mutant strain compared to the WT strain. In the mutant strain, the majority of the downregulated genes were found to be associated with the pathway of muscle development and contraction [Chrdl1 (4.16), Vgll3 (3.0), Fibin (4.6), Bmp3 (3.4), Scube2 (5.0), Inmt (4.4), Ogn (4.0), Tgfbr3 (0.88), Adra1a (5.22)] as compared to the WT strain (Figure 6C) (Supplemnatry spread sheet 3).

**Figure 6.**
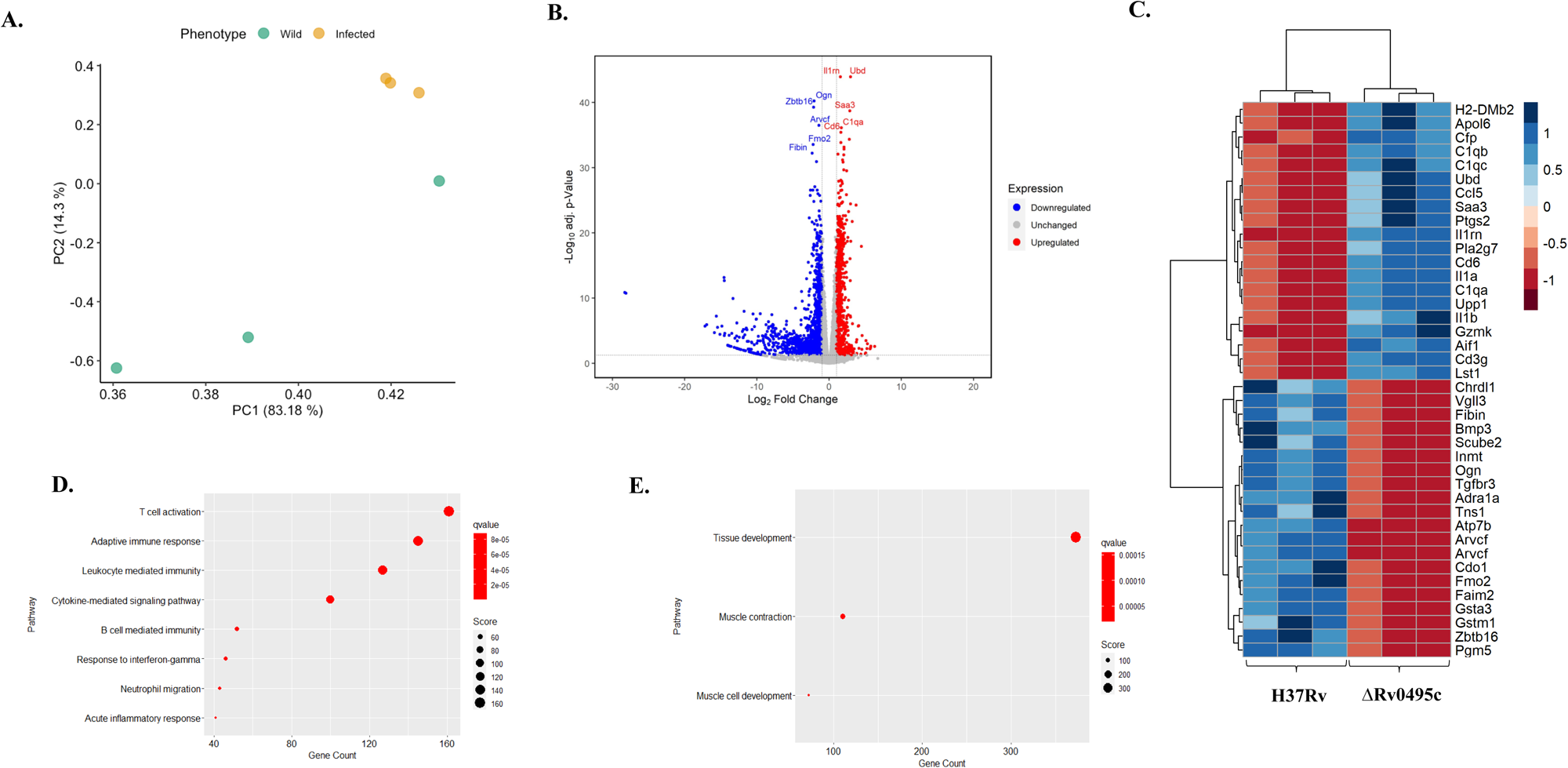
Transcriptional analysis of lung tissue from Mtb-infected mice. (A) PCA plot comparing the transcriptomes of mice lung infected with WT H37Rv and with *ΔRv0495*. **(B)** Volcano plot showing significantly deregulated protein coding genes (FDR<0.05). Blue dots indicate genes upregulates genes, grey dots represent no change whereas red dots represent downregulates genes (Diffexpressed: differentially expressed). **(C)** Heatmap showing top 20 upregulated/downregulated protein coding genes based on adjusted *p-value* (Log_2_FC > ±1). **(D)** Dotplot of positively enriched terms showing enrichment scores (*p-values*) and size of dots corresponds to gene count. **(E)** Dotplot of negatively enriched terms showing enrichment scores (*p-values*) and size of dots corresponds to gene count..

Pathway enrichment analysis of differentially expressed genes indicated that the processes related to adaptive immune response like T cell activation and B cell-mediated immunity were most significantly enriched in the *Rv0495c* mutant strain compared to the WT strain (Figure 6D). Also, the genes belonging to leukocyte-mediated immunity were increased in expression in the mice-infected with mutant strain compared to those infected by WT strain (Figure 6D). While leukocytes and the adaptive immune response play a crucial role in providing immune protection against TB, we hypothesize that the unregulated activation of these cells is primarily influenced by the significantly elevated bacterial load in the lungs of mice infected with mutant strain when compared to the WT strain. Furthermore, we observed an upregulation of genes related to neutrophil migration in the mutant strain (Figure 6D). This finding aligns with the established association between elevated bacterial load in TB and excessive neutrophil infiltration [33]. Consequently, the expression of genes related to both neutrophil migration and the acute inflammatory response was elevated in lung tissue of mice infected with *Rv0495c* mutant strain, further substantiating our findings. Simultaneously, the *Rv0495c* mutant strain infected mice lung tissue demonstrated a reduction in the expression of genes associated with tissue development and muscle development and contraction pathways (Figure 6E). This indicates its capacity to impede the recovery of a healthy lung architecture in the lungs of mice infected with the *Rv0495c* mutant strain when compared to the WT strain. Together, these data suggest that the mutant strain exhibits a hypervirulent phenotype due to excessive activation of immune cells (innate and adaptive), leading to extensive tissue necrosis and disruption of the normal tissue architecture.

## 4. Discussion

In the Mtb genome, over 40-50% of the genes are classified as conserved hypothetical with unknown function [34]. The function of these hypothetical proteins has been predicted and understood using a variety of bioinformatics tools. About 1050 hypothetical proteins were sequenced and their functions were predicted with high confidence and classified as enzymes, transporters and DNA-binding proteins [35]. However, many of these genes expressing hypothetical proteins remain uncharacterized. *Rv0495c* is one such non-essential gene that has been reported to be essential for the growth of Mtb on cholesterol[13]. In the present study, we found that the absence of the *Rv0495c* gene in Mtb perturbed the cytosolic redox balance creating an oxidized intracellular environment. Furthermore, the intracellular Fe-mediated activation of the Fenton reaction in Δ*Rv0495c* under regular growth medium resulted in an increased generation and accumulation of ROS. As a consequence, there is a signifcant decrease in the NADH:NAD^+^ ratio manifesting in the form of reduced metabolism and growth observed in the case of the mutant strain relative to the wild type [36]. The same was further validated by resazurin-based metabolic activity assay which indirectly measures cellular NADH:NAD+ ratio. The highly oxidized intracellular environment promoted lipid hydroperoxidation thereby affecting the membrane integrity of the Δ*Rv0495c*. Surprisingly, in comparison to the wild-type strain, Δ*Rv0495c* demonstrated enhanced fitness in their ability to grow inside the mouse model of Mtb infection. These above findings underscore the importance of understanding the disease biology in the context of the host and that the pathogens’ ability to utilize host machinery for its advantage is under-explored. The complete absence of the *in-vivo* growth defect observed in the absence of *Rv0495c* in mice is very intriguing. This could be attributed to the reports that suggest that Mtb is known to survive better inside the host under a mild oxidizing state [37]. Future studies focusing on characterizing these proteins would result in screening out such proteins as potential drug targets.

Our data from both the redox and transcriptional studies suggests that an enhanced level of intracellular ROS together with a lowering of the antioxidant defense system, due to a decrease in the expression of peroxiredoxin AhpC protein, created a highly oxidizing environment inside the mutant strain thereby disrupting the redox homeostasis. Disruption of the redox homeostasis affect the ability of the *Rv0495c* mutant to grow under stress which includes growth under low oxygen conditions and inside macrophages [38, 39]. We believe that the highly oxidized intracellular environment inside the Δ*Rv0495c* under regular growth medium triggered the Fenton reaction which resulted in an enhanced sensitivity to the free iron in the culture media. Similar findings have been reported by various groups whereby it has been demonstrated that a highly oxidized cytosolic environment contributed to the oxidation of cellular redox pairs such as NADH and Menaquinone, which leads to the oxidation of free iron available in the cytosol leading to ROS generation via Fenton Chemistry [40, 41].

Increased ROS could induce aberrant oxidation of biomolecules including the lipids that are essential components of the cell membrane. Our data suggest that higher levels of oxygen free radicals resulted in the degradation of the Mtb membrane lipids which resulted in destabilized cell membrane integrity of the ΔRv0495c strain under regular growth medium. The later explains the sensitivity of Δ*Rv0495c* to all anti-TB drugs tested under regular growth medium. The lower abundance of specific lipids observed in the Δ*Rv0495c* on TLC plates is very intriguing in addition to altered colony morphology. Unfortunately, the lack of mechanistic understanding limits our interpretation of this data. Both the possibilities *i.e.*, a direct or indirect role of Rv0495c in the synthesis of specific lipid molecule(s) or certain lipid molecules, due to their abundance, being more vulnerable to lipid peroxidation needs to be verified.

The reversal of the mutant phenotype in its ability to grow inside the host (*in-vivo* model) versus its ability to grow inside the macrophages (*ex-vivo* model) was very surprising. A partial rescue of the growth inside the macrophages by the addition of Fe chelators alludes to the fact that susceptibility of the free Fe in the culture media could be a reason for the growth defect observed in the *ex-vivo* model. Since it has been reported that the available Fe inside the host is always in the bound form [41] and hence the lack of free Fe could be the reason for the complete reversal of the phenotype. Although, this could not explain the enhanced growth relative to the wild type observed under *in-vivo* conditions. To gain valuable insights into the underlying host factors contributing to the distinctive phenotype observed in the mutant strain, we conducted a RNA sequencing analysis. The comprehensive data obtained from this analysis suggests that the mutant strain displays a hypervirulent phenotype as compared to the WT strain. This hypervirulence can predominantly be attributed to the uncontrolled activation of immune cells, which in turn leads to severe tissue necrosis and disruption of the normal lung tissue architecture. These findings significantly enhance our understanding of how immune cell hyperactivity contributes to the observed phenotype in the mutant strain. Again, we do not completely rule out the chances of the strain being an auxotroph of some as of yet unknown metabolite. We acknowledge that further studies focusing on elucidating the exact mechanism and role of Rv0495c protein in the pathogen’s physiology and pathogenesis will help address the above observations.

In conclusion, in this report we implicate that Rv0495c in maintaining the redox homeostasis inside Mtb. The absence of this gene in Mtb led to an increase in the abundance of intracellular ROS imparting a highly oxidizing cytosolic environment. The oxygen free radicals induce lipid hydroperoxidation mainly affecting the integrity of the lipid-rich cell membrane. This made the mutant strain more susceptible to various stress and anti-TB drugs. Finally, the observation that the absence of this gene enhances the ability of Mtb to grow better under *in-vivo* conditions opens the possibility of Mtb using this pathway to regulate its growth during infection by modulating host immune response. A better understanding will unravel the role of Rv0495c in the context of host-pathogen interaction and will open new avenues and pathways that could be targeted for developing novel intervention strategies against tuberculosis.

## Acknowledgment

Mrx1-roGFP2 redox biosensor was a kind gift from Dr. Amit Singh, IISc Bengaluru, India. This research was funded by Core funding from the Translational Health Science and Technology Institute (THSTI) to AKP, India-Singapore grants jointly sponsored by the DST in India to AKP (Project ID; INT/Sin/P-08/2015) and A*STAR in Singapore to AS (Project ID: 15302FG151). We acknowledge the support received from the IDRF (BSL3) and the Small Animal Facility (SAF) at THSTI. The sponsors played no part in its planning, gathering, analyzing, or interpreting data, or in the preparation of the report. The corresponding author had full access to all the data in the study and had final responsibility for the decision to submit for publication.

## 5. Author contributions

AKP and RP contributed to the conception and design of the study. RP, STa, MP, VN, TS, STy, VB were responsible for the acquisition of data. RP, STa, AKP analyzed and interpreted the data. SC and RN performed the ICP-MS and analyzed the Host RNAseq data. SKG and YK performed and analyzed the metabolomics data. AS generated and analyzed the bacteria RNAseq data. RP and AKP drafted the manuscript and all authors were involved in revising it critically for intellectual content and have given final approval of the version to be published.

## Supplementary Figures

**Figure S1.** level of NADH/NAD^+^ reduction was also measured using MTT kit at 6^th^ and 12^th^ day of growth curve when started at OD_600_ 0.01. data was normalized with CFU at their respective days.

**Figure S2.** Estimation of intracellular ROS generation schematic figure depicting the response of DCFH-DA with ROS in fluorescence generation whereas the graph represents mean fluorescent intensity (MFI) graph of *H37Rv, ΔRv0495c*, and *ΔRv0495c:0495c* strains using CHP as a positive control and ascorbic acid as a negative control as described in materials and methods.

**Figure S3.** Intracellular iron levels of *H37Rv, ΔRv0495c*, and *ΔRv0495c:0495c* Mtb strain grown in 7H9 medium enriched with 10% OADS and 0.05% Tween 80 was measured at 6^th^ day using ICP-MS. *y-axis* depicts the concentration of iron in ppb/bacteria with respect to time (days) on x-axis. Data represented here was normalized with CFU, and replicated in *n = 3* biological and averages for two independent sets of experiment.

**Figure S4.** *H37Rv, ΔRv0495c* and *ΔRv0495c:0495c* strains were spotted on 7H11 agar plates supplemented with OADS and incubated for 21 days at 37^0^C and the spotted colonies were photographed.

**Figure S5. (A)** 1D-TLC analyzed using solvent [Chloroform: Methanol: water (65:25:4, v/v/v)] for the analysis of phospholipids (PIMs, PE, CL, PI, Acylated-PIMs) and developed by iodine fumes. **(A)** 2D-TLC analyzed using solvent [Chloroform: Methanol: water (60:30:6, v/v/v)] for 1D and for 2D TLC was run using Chloroform: Methanol: Acetic acid: Water (40:3:25:6 v/v/v/v) plate was then dried and visualized by iodine vapors.

## Graphical Abstract

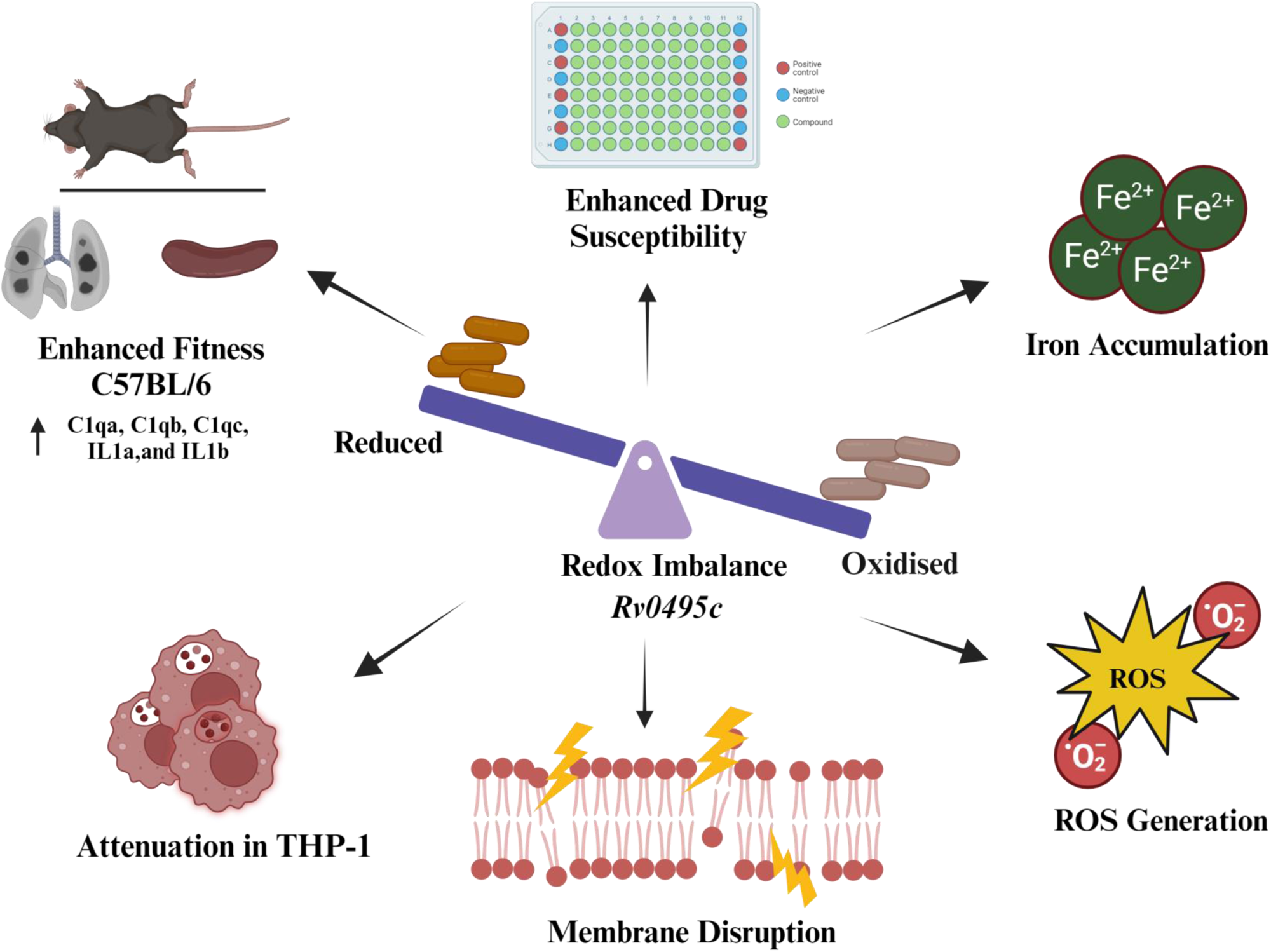

