## Supplementary figures and images for "Hypothetical gene *Rv0495c* regulates redox homeostasis in *Mycobacterium tuberculosis*"

### Supplementary Figures

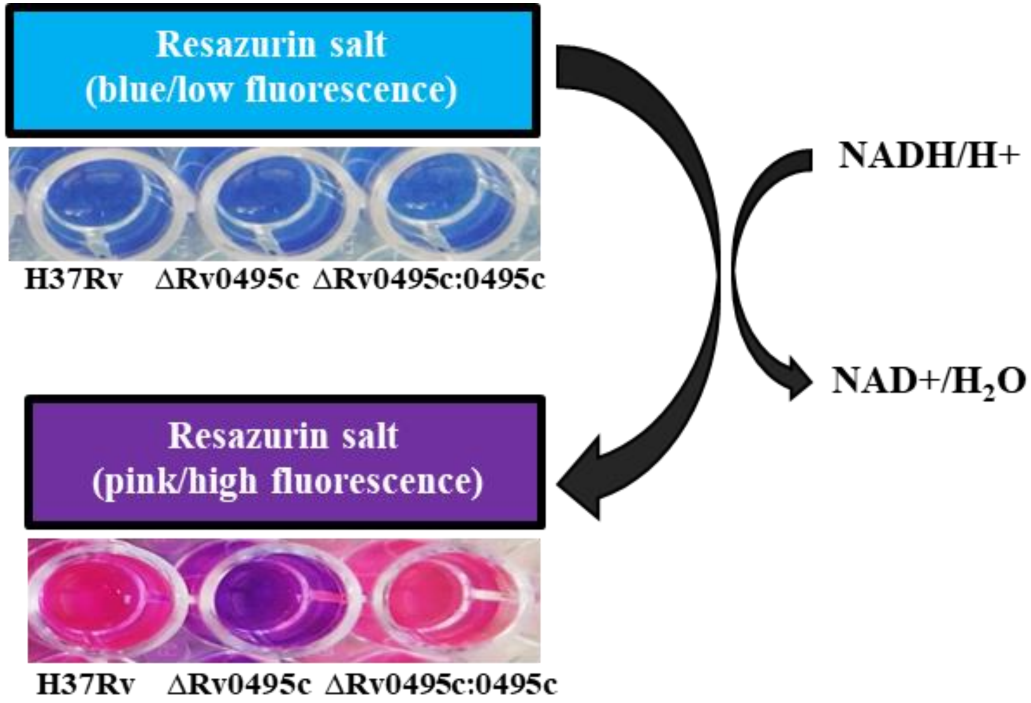

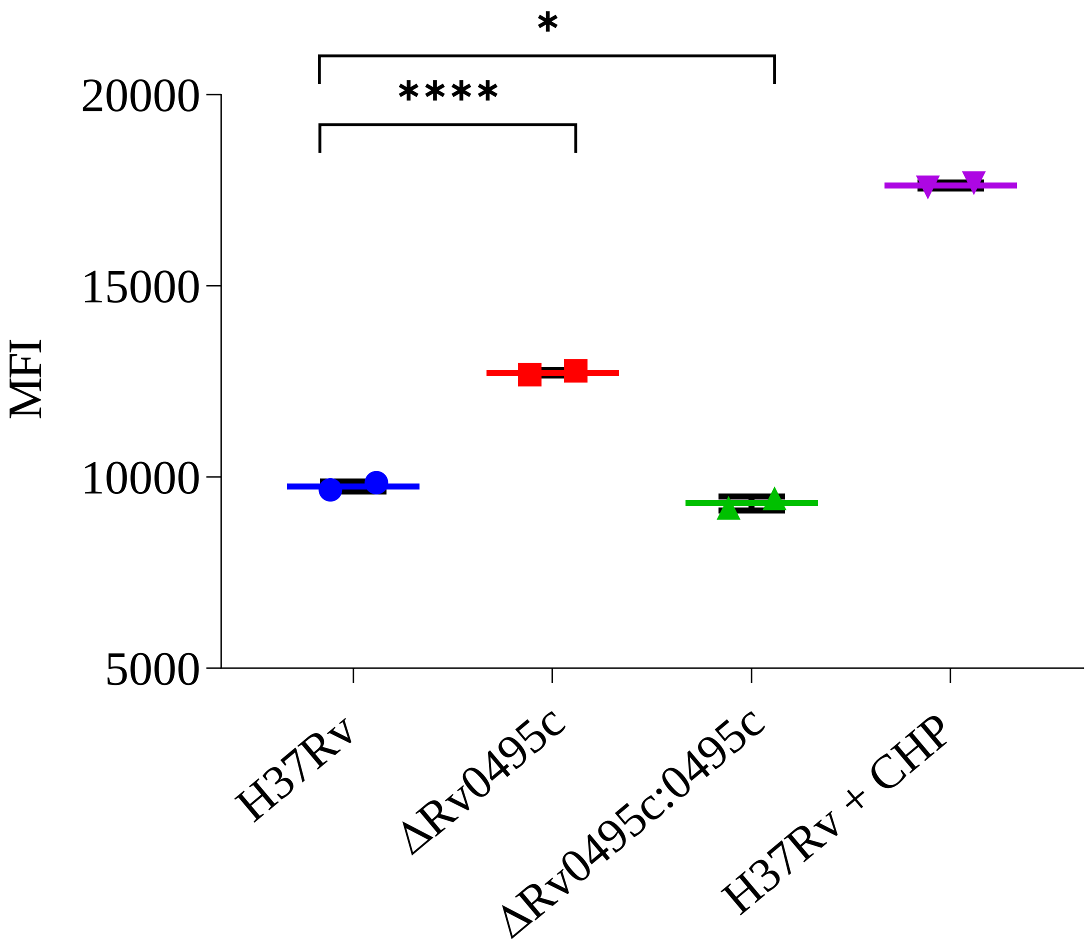

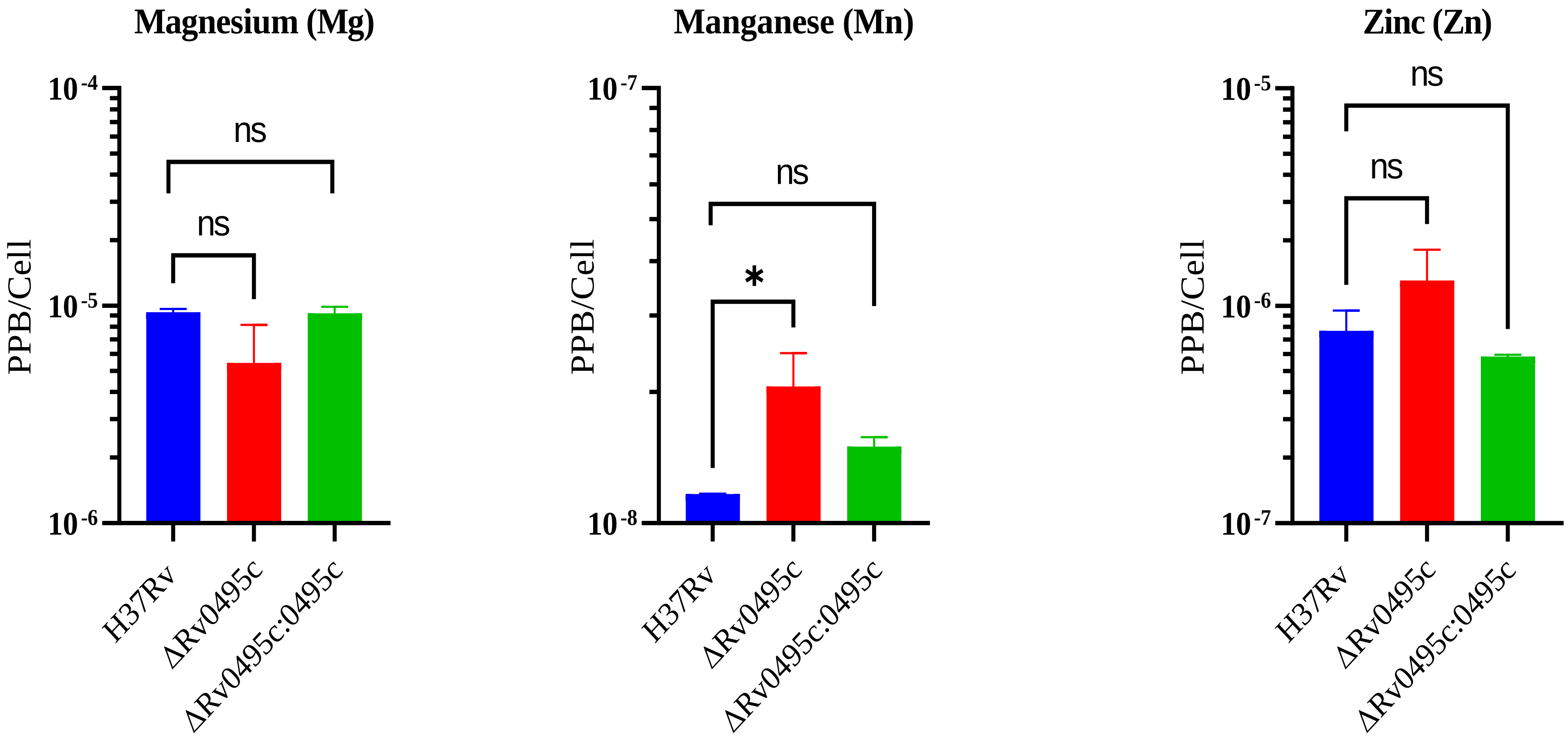

**Figure** S4.

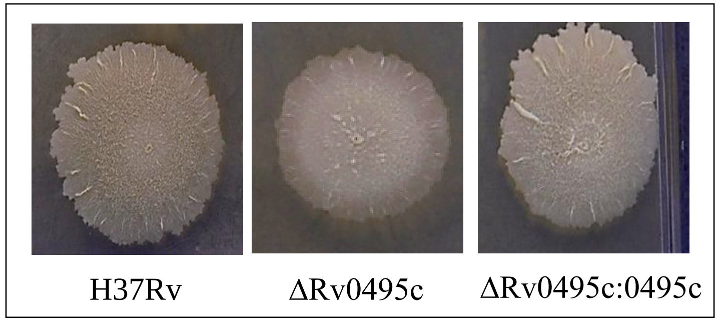

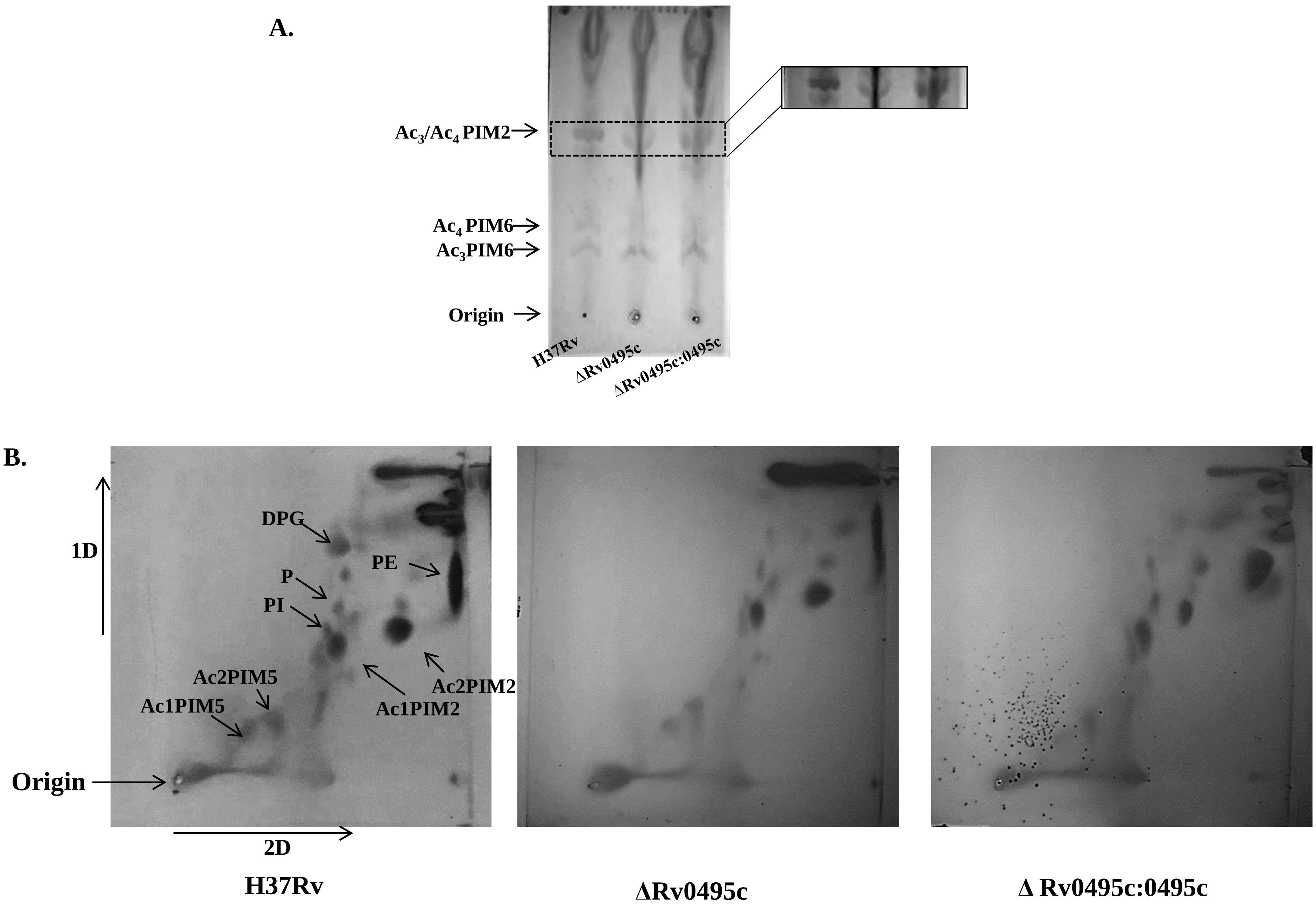
