## Supplementary Tables for "Hypothetical gene *Rv0495c* regulates redox homeostasis in *Mycobacterium tuberculosis*"

**Table S1. List of primers used in this study**

| **S.no** | **Primers** | **Sequence (5’-3’)** |
| --- | --- | --- |
| 1. | Rv0495-F1 | ATGATATCGGGCAGATTTCCACCGCTTCTAT |
| 2. | Rv0495-R1 | ATGCGGCCGCGGATTCAACGTTAGACCACGAAGC |
| 3. | Rv0495-F2 | ATCTCGAGAGTCAATTAGGGCTCATCGCCG |
| 4. | Rv0495-R2 | ATCTCGAGCGTCTCACAGTGCACCCATAG |
| 5. | 0495-conf1 | TTCCGTCGGCTTCGCTGTC |
| 6. | 0495-conf2 | GTGGAAACCAGGTCACCCAG |
| 7. | 0495-F | GTTCAGCTGTTAATTAAATACGGGCGTGTGGCGAC |
| 8. | 0495-R | CAGATTTAAATTTAGCTACTGGGCCGCACG |

**Table S2. List of strains used in this study**

| S. No | Strains/plasmids | Description | Source |
| --- | --- | --- | --- |
| 1. | XL-1 Blue (*E. coli*) | *recA1 endA1 gyrA96 thi1hsdR17 supE44 relA1 lac [F’ proAB lacIq Z∆M15 Tn (Tet^r^)]* | Stratagene, USA |
| 2. | H37Rv | *Mycobacterium tuberculosis* Parental Strain | A generous gift from Christopher M. Sassetti. |
| 3. | *∆Rv0495c* marked | Deletion mutant strain of H37Rv lacking *Rv0495c*, harboring Hyg^r^ | This Study |
| 4. | *∆Rv0495c* unmarked | Unmarked mutant strain of H37Rv lacking *Rv0495c* | This Study |
| 5. | ∆*Rv0495c:0495c* | Complemented Strain, *Rv0495c* integrated at L5 *attB* site in ∆*Rv0495c* mutant. | This Study |
| 6. | pJM1 | Suicidal vector for generation of Allelic Exchange Substrate (AES) | A generous gift from Christopher M. Sassetti. |
| 7. | pJEB402 | Shuttle Expression vector, Kan^r^, Integrative. | A generous gift from Christopher M. Sassetti. |
| 8. | pMV762:Mrx1-roGFP2 | Redox biosensor, Hyg^r^ | *Bhaskar A.et.al., 2014* |
| 9. | Rv:Mrx1-roGFP2 | Wild type Rv, expressing redox biosensor | This study |
| 10. | ∆Rv0495c:Mrx1-roGFP2 | *Rv0495c* null mutant expressing redox biosensor | This study |

**Table S3. Solvent System used for TLC analysis**

| **Lipid Class** | **Solvent system** |
| --- | --- |
| Mycolic Acid | Hexane: ethyl acetate (95:5 v/v) |
| Phospholipids | Chloroform: Methanol: Water (65:25:4 v/v/v) |
| Apolar lipid  (PE, PI, PIMs) | 1-D Chloroform: Methanol: Water (60:30:6, v/v/v)  2-D Chloroform: Methanol: Acetic acid: Water (40:3:25:6 v/v/v/v) |
